# Fully functional AAV viral vectors with highly altered structural cores and subunit interfaces using ProteinMPNN

**DOI:** 10.1101/2025.07.24.666527

**Authors:** Ziyu Jiang, Sirimar Laosinwattana, Paul A. Dalby

## Abstract

Adeno-associated viruses (AAV) have emerged as a viable vector for gene therapy, with several clinical approvals and a growing pipeline in clinical trials. These vectors have several challenges that need to be addressed to widen their use, including improving tropisms, reducing manufacturing costs, increasing storage stability, minimising their immunogenicity, or evasion of existing AAV immunity in which neutralising antibodies lead to loss of potency. Vector engineering, particularly capsid protein engineering, offers a potential route to generating new capsids that can selectively target desired cell types or evade pre-existing immunity, while also ensuring that they are manufacturable at higher titres, more stable, and have reduced immunogenicity. Extensive protein redesign is emerging as a viable option, through generative AI approaches, for engineering many types of protein. Here we explored the potential of ProteinMPNN, to extensively redesign AAV2 and yet still form stable and functional capsids. We targeted 52% of AAV2 residues for redesign by ProteinMPNN, such that only the buried protein core and subunit interfaces would be varied, leaving the capsid external and internal surface features unchanged. The aim was to significantly modify the structure responsible for assembly and capsid integrity, while maximising the probability of maintaining the wild-type DNA packaging and transduction capabilities. The final designs were between 14% and 30% mutated overall, and yet were capable of forming functional and intact capsids, with the transduction efficiency of wild type retained for some variants. The designs generally led to lower titres from cell culture, yet some designs had either improved capsid packaging efficiency or transduction efficiency. In particular, our “Pentamer” design had the best transduction efficiency, while our “Chimera” design had a packaging efficiency that was 2.5x higher than for the WT AAV2. These results demonstrate the potential to use generative AI tools in vector capsid redesign for novel core assembly features, and now pave the way for expanding this approach into selectively re-engineering their surface properties to influence tropism, immunogenicity and transduction efficiency.

## Introduction

The rapidly developing field of protein engineering has emerged as a critical area of research. Adeno-associated virus (AAV), a small single-stranded DNA virus with an icosahedral protein capsid structure, has been extensively used in gene therapy developments, owing to their low pathogenicity and broad infection tropism (Wang *et al*., 2024). AAV capsids are comprised of 60 protein subunits formed into an icosahedral structure. The subunits come from three viral proteins, virion protein 1, 2 and 3 (VP1-3), at a ratio of typically 1:1:10, whereby the capsid structure itself is assembled through the VP3 structure that is common to all three VPs (Xie *et al*., 2002).

The wild-type AAV genome comprises two open reading frames (ORFs), *rep* and *cap* flanked by two inverted terminal repeats (ITRs). The *rep* gene encodes for proteins that are essential for viral replication assembly (Rep78, Rep68, Rep52, and Rep40), whereas the *cap* gene encodes for VP1-3 (Collaco *et al*., 1999). To enhance the safety of therapeutic adeno-associated viral vectors, recombinant AAVs (rAAVs) were designed with the *rep* and *cap* genes removed from the genome, and so requiring multi-plasmid systems for production in mammalian cell lines. The first rAAV system was produced by transfecting helper adenovirus (Ad2)-infected human D6 cells with two plasmid constructs. The first replaced the wild-type AAV genome with a neomycin-resistance gene (*neo*), and the second contained the *rep* and *cap* genes (Hermonat & Muzyczka, 1984). The successful transduction of the neomycin resistance gene into mammalian cells marked the foundation of the development of rAAV-based gene therapy. Removing the wild-type AAV *rep* and *cap* genes from the vector genome, the resulting rAAVs were replication-deficient, which significantly improved their safety as therapeutic vectors (Suarez-Amaran *et al*., 2025). Nevertheless, the use of adenovirus-infected producer cells retained the risk of generating replication-competent, wild-type-like rAAVs (rcAAVs) (Allen *et al*., 1997). To address this, helper-free rAAV packaging systems were developed with a third plasmid carrying the adenoviral helper functions (*E2A, VAI* and *E4* genes) (Collaco *et al*., 1999). These advances in safety and the resulting improved productivity of rAAV enabled the successful launch of various rAAV-based gene therapy products, such as Luxturna (Spark Therapeutics, Inc), Glybera (uniQure biopharma B.V.) and Zolgensma (Novartis AG).

Despite the early successes, the broader development of AAV-based gene therapy products remains limited by their cell tropism, inherent capsid instability, especially for AAV2, and the presence of pre-existing immunity to wild-type AAV in some populations Jiang and Dalby, 2023; Murphy *et al*., 2008, Manno *et al*., 2006). Capsid protein engineering efforts have primarily aimed to modify the tropism of AAVs, with some variants also having increased thermal stability (Santiago-Ortiz *et al*., 2015; Bennett *et al*., 2023). The conventional and commonly used techniques, such as directed evolution, consensus mutations, and ancestral sequence reconstruction, while successful (Santiago-Ortiz *et al*., 2015; Liu *et al*., 2021; El Andari *et al*., 2022; Rode *et al*., 2022), can be time-consuming and limited by selection *in vitro* or in mouse models, and also cannot easily obtain significantly altered yet functional protein sequences that would aid in designing vectors with no pre-existing immunity.

Recent machine learning and deep learning technologies have revolutionised the field of protein engineering. Deep-learning protein sequence design tools such as ProteinMPNN (Dauparas *et al*., 2022), RFdiffusion (Watson *et al*., 2023), ProteinGAN (Repecka *et al*., 2021) and ABACUS-R (Liu *et al*., 2022) have gained rapid and remarkable success in the design of *de novo* protein structures. ProteinMPNN is a deep-learning-based protein sequence design method that harnesses the power of neural networks to predict amino acids that fold into the desired protein structure (Dauparas *et al*., 2022). By using a structure-based algorithm, ProteinMPNN is able to generate protein sequences that fold more confidenly into the input structure than the native sequence, suggesting the potential of ProteinMPNN to improve protein stability (Dauparas *et al*., 2022). However, due to the solely structure-based method, design constraints and intensive screening efforts are often still required to select candidates with retained functionality (Sumida *et al*., 2024).

In this study, ProteinMPNN was used to design novel adeno-associated virus capsids with the aim of modifying a significant proportion of the wild type sequence. This would potentially offer a route to generating vectors that are not recognised by existing neutralising antibodies. It also explored the extent to which largely buried and subunit interface residues could be redesigned while retaining wild-type like AAV2 structure and function. To further explore the sequence space accessed by the ProteinMPNN designed variants, ancestral sequence reconstruction (ASR) was applied to generate ’virtual ancestors’ based on a multiple sequence alignment and similarity-based clustering of the ProteinMPNN-generated sequences. The purpose of these virtual ancestors was not to recreate true evolutionary intermediates, but purely as a method to generate additional potentially viable hybrid sequences from the designed variants. The structure and function of these AAV capsid variants were subsequently evaluated to demonstrate the potential of machine learning-guided approaches for extensive AAV redesign.

## Methods

### Solvent Accessible Surface Area Analysis

To be able to place constraints on the exposed residues, a Solvent Accessible Area Analysis (SASA) was performed using BIOVIA Discovery Studio. The wild-type AAV2 monomer PDB file was loaded into Discovery Studio. SASA analysis was carried out using the default setting where residues with more than 25% solvent accessible surface (SAS) were considered ‘exposed’ and residues with less than 10% solvent accessible surface (SAS) were considered ‘buried’. The results produced from the SASA analysis contain the SAS percentage of each residue. The hydrophobicity score of each residue was also given, the more positive the score, the more hydrophobic the residue is and the more negative the score, the more hydrophilic the residue is.

### ProteinMPNN-guided capsid design

An AAV2 capsid structure (PDB ID: 1LP3) was fetched from RCSB Protein Data Bank (PDB). The fivefold and threefold interface pentamer and trimer of protein subunits were selected from the fetched PDB file using Pymol. The pentamer and trimer of proteins were then exported as individual objects, producing separate PDB files. To design novel AAV sequences, the pentamer and trimer PDB files were input into the ProteinMPNN huggingface interface (Dauparas *et al*., 2022). The sampling temperature was set to 0.1 to allow maximum sequence recovery. All highly exposed surface residues were fixed to retain the solubility and function of the output sequences. 150 pentamer and 150 trimer sequences were generated and aligned using Clustal Omega (EMBL-EBI) (Madeira *et al*., 2024). A consensus sequence was produced for each of the two multiple sequence alignments based on the percentage of conservation within the alignment. Only residues with at least 90% conservation were included in the consensus, otherwise the original wild-type residue was retained.

### Designed capsid structure screening

The consensus sequences were folded using AlphaFold2 (Jumper *et al*., 2021) to provide a preliminary estimation of the foldability of the protein sequence and to produce PDB structure model files. Template modelling scores (TM-score) between the wild-type AAV2 monomer structure and the AlphaFold2-predicted structure were calculated using a pairwise alignment tool TM-align (https://zhanggroup.org/TM-align/) (Zhang and Skolnick, 2005). To re-create a protein design that contained both of the newly designed threefold and fivefold interfaces in the AAV capsid, the trimer and pentamer sequences generated by ProteinMPNN were chimerised manually, taking their respective interface residues. The output chimaera sequence was then input back into ProteinMPNN with constraints placed on the residues at the fivefold and threefold protein interfaces, together with the exposed surface residues. Therefore, only core residues that were not located at the subunit interfaces or capsid surface were allowed to mutate.

### Virtual ancestor chimera sequence reconstruction

From the 150 ProteinMPNN-designed chimeric sequences, a phylogenetic tree and a multiple sequence alignment (MSA) were produced using Clustal Omega, exported into FASTA format (Madeira *et al*., 2024), and the phylogenetic tree (constructed with the default Unweighted Pair Group Method with Arithmetic Mean method of Clustal Omega) was exported into a Newick file. These two files were then put into a Graphical representation of ancestral sequence reconstructions (GRASP) (Foley *et al*., 2022), to generate 150 (numbered from 0 to 149) virtual ancestor sequences of the ProteinMPNN-designed chimera sequences.

### In silico solubility screening

Solubility screening was performed by uploading the Fasta file containing all sequences into the online CamSol portal (Sormanni, Aprile and Vendruscolo, 2015; Sormanni *et al*., 2017),where the intrinsic solubility of each sequence was predicted.

### AAV production

HEK293T cells were seeded at a density of 25000 cells/cm^2^ in T25 flasks and grown in Dulbecco’s Modified Eagle Medium (DMEM) (Cat. No. 11965092, ThermoFisher Scientific) with 10% Fetal Bovine Serum (FBS) and cultured at 37 °C. Cells were expanded in T75 flasks 48 to 72 hours post-seeding and further expanded in T175 flasks 48 to 72 hours after expansion in T75 flasks. To ensure optimum production titre, at 90% confluency, HEK293T cells were further expanded in three-layer flasks with ventilated caps. Inspection of the cells under the microscope was performed daily during culture to determine cell confluency. At 70% to 80% confluency cells were transfected with 68 ng cm^-2^ AAV2 packaging plasmid (pAAV2-2), 52 ng/cm^-2^ green fluorescence plasmid (pAAV-CAG-GFP), and 146 ng cm^-2^ helper plasmid (pAdDeltaF6) using linear polyethylamine (PEI) to initiate viral vector production.

### AAV purification

The harvested cells were lysed via three cycles of freeze-thaw and incubated with 50 U/ml benzonase (Cat. No. E8263, Sigma-Aldrich) for 45 minutes at 37 °C. The cell debris in the lysate was removed by centrifugation, and the supernatant was collected and filtered through a 0.45 µm syringe filter (Cat. No. SLHPR33RB, MERK). To produce a concentrated virus stock, the filtrate was added to the 100 kDa spin column and spun at 2250 g in a bench-top centrifuge. This spin-down process was repeated until the volume of the lysate is concentrated to ≤ 1.5 ml. The concentrated lysate was then transferred to autoclaved 1.5 ml Eppendorf tubes and stored at −80 °C Viral vectors were purified via iodixanol gradient ultracentrifugation. The iodixanol gradient (60%, 40%, 25%, 15%) was prepared from OptiPrep Density Gradient Medium with 60% iodixanol (Cat. No. ab286850, Abcam) and PBS-MK (Phosphate-buffered Saline with MgCl2 and KCl). 5 ml of 60% iodixanol was first added to the bottom of the ultracentrifuge tube, followed by overlaying of the 40% iodixanol, 25% iodixanol and then the 15% iodixanol. Sample lysates were loaded on top of the 15% iodixanol and topped up with PBS to fill the ultracentrifuge tube, followed by ultracentrifugation at 274,000 rpm for 5 hours and 15 minutes. After ultracentrifugation, purified samples were extracted from the 25% iodixanol layer, and buffer exchanged into PBS-MK using Vivaspin® 20 Centrifugal Concentrator (Cat. No. VS2041, Sartorius) to remove iodixanol.

### AAV particle quantification

AAV2 Xpress ELISA kit (Cat No.PRAAV2XP, Progen) was used to quantify the total viral capsid present in purified AAV samples by following the manufacturer’s protocol. Each sample was diluted 20 times with ASSB 1X buffer prior to the start of the experiment.

### dPCR AAV genome copy quantification

dPCR quantification on the AAV genome was performed by Mr Ben Jones at Queen Mary Blizard Institute using the SYBR Green technology protocol. Samples were treated first with DNase I for 30 minutes at 37 ℃ and 95 ℃ for 15 minutes, followed by proteinase K for 30 minutes at 55 °C, 15 minutes at 95 °C and 5 minutes at 4 °C.

### AAV transduction analysis through flow cytometry

0.01×10^6 HEK293T cells were seeded in 96 wells, and on top of the cells, 55 µL of purified AAV samples were added. For each sample, biological triplicates were made. 48 hours post-transduction, cells were harvested by first removing the spent media and then aspirate with PBS. Re-suspended cells were analysed by using the LSRFortessa cytometer and the results obtained were analysed using FlowJo 10.8.

### Transmission electron microscopy imaging of AAV particle structure

TEM images were acquired using the negative staining methods by Mr Shu Chen at Birbeck, University of London. 1 µL of purified sample was used for each imaging experiment. Samples were first reloaded onto the TEM support copper grid, followed by air-drying for 5 minutes and staining with 2% uranyl acetate for 30 seconds prior to imaging.

## Results and Discussion

### Design strategy

The design strategy for AAV capsids was based on the capsid structure for AAV2, comprised of 60 subunits (Figure 1A) at a ratio of approximately 1:1:10 for VP1 : VP2 : VP3, and formed into a T=1 icosahedron (Wistuba *et al*., 1995; Sonntag, Schmidt and Kleinschmidt, 2010; Aydemir *et al*., 2016; Wörner *et al*., 2021). However, VP1 and VP2 all contain the same sequence and structure of VP3 (PDB: 1LP3) as all three subunits are derived from the same gene. The capsid assembly itself is determined entirely through the structure and subunit interactions at the 5-fold, 3-fold and 2-fold interfaces, and involve only the VP3 region that is common to all subunits.

**Figure 1.**
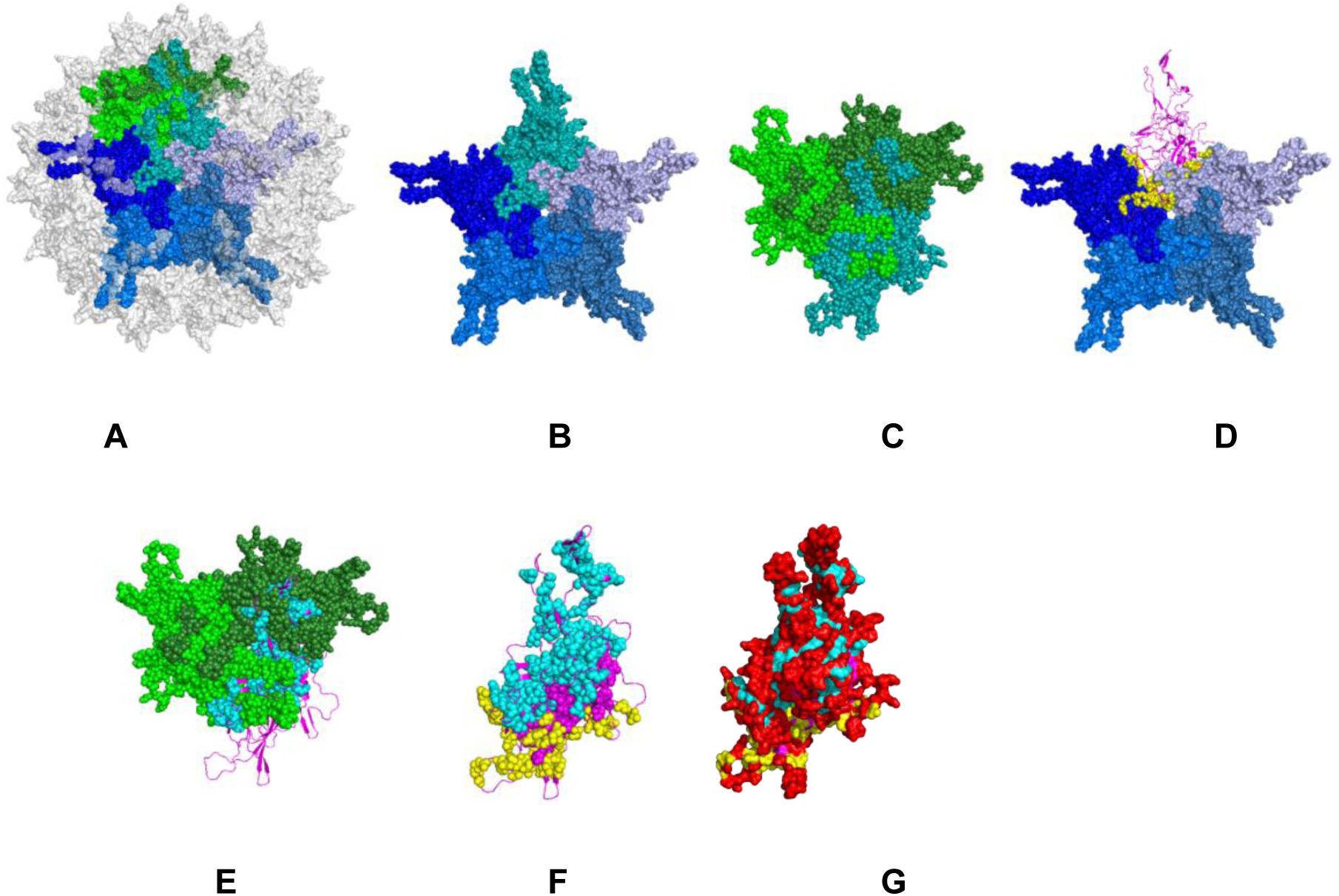
AAV capsid design strategy. A) whole capsid of 60 VP3 subunits highlighting one pentamer (blues) and one trimer (greens), with their common monomer shown as teal spheres. This monomer is highlighted in the B) pentamer as teal spheres, C) trimer as teal spheres, D) pentamer as pink ribbons with pentamer interface residues highlighted as yellow spheres, and E) trimer as pink ribbons with trimer interface residues shown as cyan spheres. F) VP3 monomer with the 52% mutatable residues shown as spheres, coloured yellow (pentamer interface), cyan (trimer interface) or magenta (core). G) VP3 monomer in surface representation with the unmutated residues (48%) highlighted in red. Images are based on PDB ID: 1LP3 and generated in Pymol (Schrödinger LLC, 2015).

ProteinMPNN is designed to repack the amino acid side chains to maximise the probability of containing structural elements that can be combined to fold and stably form the larger target input structure, as trained by the structures available in the protein data bank (PDB). For a protein subunit such as VP3, it would be challenging to use ProteinMPNN directly on the whole capsid assembled with 60 subunits. However, a complete redesign of an isolated monomer of VP3 using ProteinMPNN, with no context of the whole capsid assembly provided, would also be expected to lose the critical subunit interactions at the 2-fold, 3-fold and 5-fold axis interfaces, required for assembling and stabilising capsids in the correct icosahedral structure. It would also be expected to lose solvent-exposed surface features that are important for cell receptor or heparin sulphate proteoglycan binding, internal interactions with packaged DNA, and potentially any important interactions with accessory proteins involved in capsid assembly and DNA packaging. Our strategy was therefore to redesign buried core and interfacial sequence regions, while retaining the solvent-exposed sequence of the wild-type AAV2 parent, with the aim of maintaining the existing capsid functions of DNA packaging, cell recognition and transduction.

To enable the subunit interfaces to be redesigned, as well as the buried core protein residues, two parallel intermediate designs were generated using structures of only the 5-fold axis pentamer (Figure 1B) and 3-fold axis trimer (Figure 1C) as inputs for ProteinMPNN. In both routes, only residues in the protein core and all subunit interfaces (Figures 1 D-F) were allowed to be redesigned, thus increasing the probability that the final capsid would retain the cell binding, DNA packaging and accessory protein interaction functionalities of the parent AAV2. In total, 267 residues (52%) were allowed to mutate (Figure 1F and Supplementary information Figure S1), leaving 48% unmutated (Figure 1G) at the highly solvent-exposed surfaces of the intact capsid. The 2-fold axis was not independently optimised with a dimeric structure as it only has a relatively minor contribution to the total subunit interface, with just 25 residues involved. The same 25 residues from each subunit interact with each other at a dimeric interface with a 2-fold rotational symmetry. Of the 25 residues at the 2-fold interface, only 7 were allowed to mutate in ProteinMPNN.

ProteinMPNN generated 150 sequences for each of the pentamer and trimer structures, and the consensus sequence of each was selected, retaining mutations only where they were at least 90% conserved across all designs. The final pentamer sequence design (Figure S1e) mutated 155 (30%) residues, while the trimer sequence design (Figure S1f) mutated 71 (14%) residues. For the trimer design, the five-fold axis interface was allowed to mutate unconstrained by any interfacial subunits, and so assembly of the five-fold axis interface to generate a full capsid was not guaranteed. The same was true for the three-fold interface that was unconstrained when redesigning VP3 within the pentamer model. To generate a VP3 structure in which both interfaces were redesigned in the presence of their respective neighbouring subunits, an initial chimera sequence was manually created from the trimer and pentamer structure redesigns, which combined the three-fold axis interface mutations from the trimer and the five-fold interface mutations from the pentamer. The initial chimera structure was predicted using Alphafold 2, and then fed back into ProteinMPNN as a monomeric protein, such that only the buried core residues that were not located at either of the subunit interfaces, or on the capsid surface were allowed to mutate again (magenta spheres in Figure 1F). Thus, the core structure was re-optimised to accommodate each of the two new subunit interfaces, while still leaving the capsid surface properties unaltered from the parent AAV2.

The final chimera design was the consensus sequence from 150 ProteinMPNN output sequences, and retained almost all of the core mutations already found, mainly in the pentamer design, to give a total of 131 mutations (25% of the total VP3 sequence). Interestingly, half of the sites that were free to mutate in ProteinMPNN retained the WT AAV2 residue. This was at least in part due to a bias away from mutating the more conserved residues in known AAV sequences, with a median conservation score of 0.38 for mutated sites, and 0.49 for sites that retained the wild-type residue, compared to an overall median score of 0.39 for all residues. Interestingly, within the sites that were mutated, there was no correlation between the conservation scores of residues and the degree of similarity for the mutation made, according to BLOSUM62. In other words, mutations at highly conserved sites were not biased towards more conservative mutations. In addition, the mutations observed were often not mutations found in natural sequences at all. Of the 131 total mutations, 101 were to mutations observed in <10% of natural sequences, and 29 were to mutations not previously observed in nature. Very low BLOSUM scores (−3, −4) occurred at 19 sites. These ranged from low to very highly conserved, including five of the seven most conserved sites mutated (>79% conserved), while the specific mutations were rarely or never seen in natural sequences. Only 17 mutations were observed in over 20% of natural sequences. This highlights the potential of ProteinMPNN to generate mutations that would normally be considered as very likely to disrupt structure and/or function. This deviation from natural sequences also highlights the potential of using ProteinMPNN to design new AAV capsids that can erase certain features such as the epitopes for neutralising antibodies, that have resulted from exposure to natural AAVs or other AAV-based gene therapies. Such an advance would have a major impact on widening the reach of AAV-based therapies to more diseases and patient numbers. Of course, the current analysis is based on the mutation of sites that are mainly buried, and it is yet to be seen whether the same exploration of new mutations is possible on the surface while retaining certain functions.

### Location of mutations

The distribution of final mutations made in the chimera consensus design is shown within the structure in Figure 2, and by sequence in Figure S1g (supplementary information). There was no obvious bias in the distribution of final mutations compared to the allowable mutation sites, in terms of their location within the structure or sequence. The final distribution of mutations between the pentamer interface (27 mutations, 20.6%), trimer interface (49 mutations, 37.4%) and core residues (55 mutations, 42%), was also essentially unchanged from the input mutable residue distributions (18% pentamer interface, 46% trimer interface, 36% core), aside from a slight bias away from trimer interface in favour of core mutations. Of the 7 residues allowed to mutate in ProteinMPNN at the 2-fold interface, only two were mutated in the final trimer design, five in the pentamer design, and just three in the final consensus chimera.

**Figure 2.**
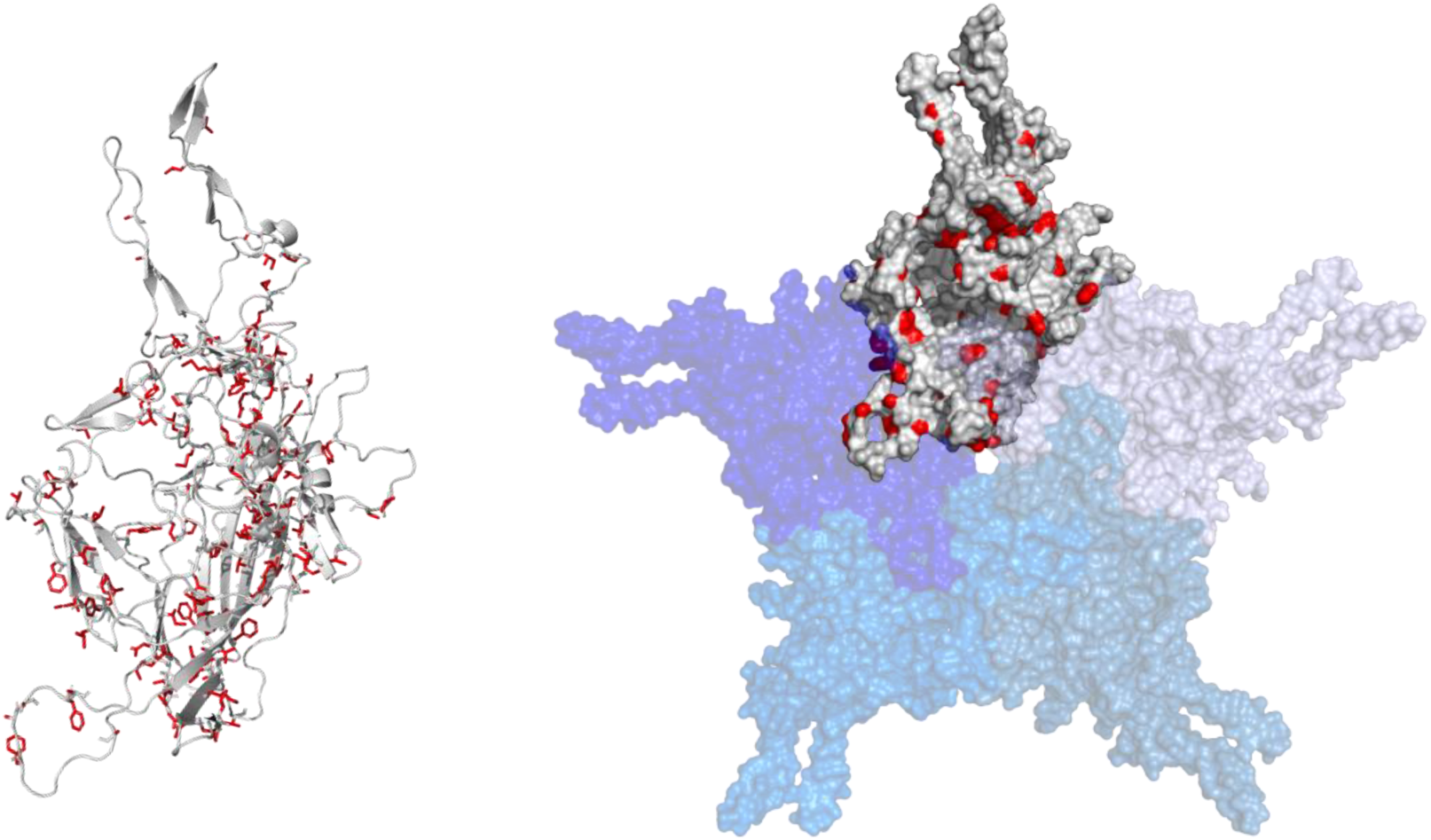
Mutation sites in chimera design. A) WT AAV2 monomer with mutated sidechains shown in red sticks. B) WT AAV2 monomer (grey) within the pentamer, showing surface and mutated sidechains in red. Images are based on PDB ID: 1LP3 and generated in Pymol.

### Virtual ancestral reconstruction and solubility screening

While the above approaches relied on using the consensus sequence from 150 outputs from ProteinMPNN, it was unclear whether this would be the best option. Therefore, given that the sequences were highly related, we used ancestral sequence reconstruction (ASR) methods on the output sequences to generate alternative sequences that further explored the overall output sequence space. It is a technique that deduces protein/DNA sequences from multiple sequence alignments and reconstructed phylogeny of input homologous sequences (Thornton, 2004). Such methods are often used to generate protein sequences that lead to functional protein structures that are more thermostable than their extant descendants (Thomson, Carrera-Pacheco and Gillam, 2022), but it was unknown whether its application to ProteinMPNN outputs would have a similar outcome. The conventional ASR method involves the alignment of homologous sequences and inferring phylogeny. Second, the ancestral state of each sequence site is determined, and the predicted ancestral sequences are generated physically. Finally, the expressed ancestral sequences are screened for the desired functional properties (Thornton, 2004). The 150 new output sequences generated in ProteinMPNN for the chimera were therefore further analysed as if they were evolutionarily related sequences to construct a phylogeny (Figure S2, supplementary information) and so determine virtual ancestral sequences as additional potential candidates.

The selection of the virtual ancestor chimera sequences to take into experimental testing was based solely on their solubility scores as predicted by Camsol (Sormanni, Aprile and Vendruscolo, 2015). The solubility scores for all ancestors can be seen as blue dots in Figure S3 (Supplementary information), compared to that of wild-type AAV2 VP3 shown at position X=1 as a red dot. Essentially all virtual ancestors had predicted solubility scores significantly higher than the wild-type AAV2 VP3 sequence, suggesting the general capability of ProteinMPNN to design highly soluble sequences. Ancestors 148 and 149 (named N148 and N149, respectively) were the two ancestors with the highest predicted solubility scores, and so were also selected for experimental testing, together with the consensus chimera, and the consensus from each of the pentamer and trimer sequence designs. Compared to the chimera sequences directly output by ProteinMPNN, the virtual ancestors show an overall more compact distribution of solubility scores, with the majority harbouring solubility scores between +0.6 and +0.8. While the ProteinMPNN output sequences show a more scattered distribution of solubility scores between +0.4 to +0.8.

### Structural Alignment and TM-score between wildtype AAV2 monomer and Chimera, Ancestor N149, N148

To ensure the structural similarity between the variants and the wild-type AAV, the sequences of variants N149, N148 and the final consensus Chimera were input into AlphaFold 2 to generate predicted VP3 monomer protein structures. The structures were then assessed using the template modelling score (TM-Score) for their similarities to the wild-type AAV2 VP3 monomer. As shown in Table 1, all variants showed high structural similarities to wild-type

**Table 1.**
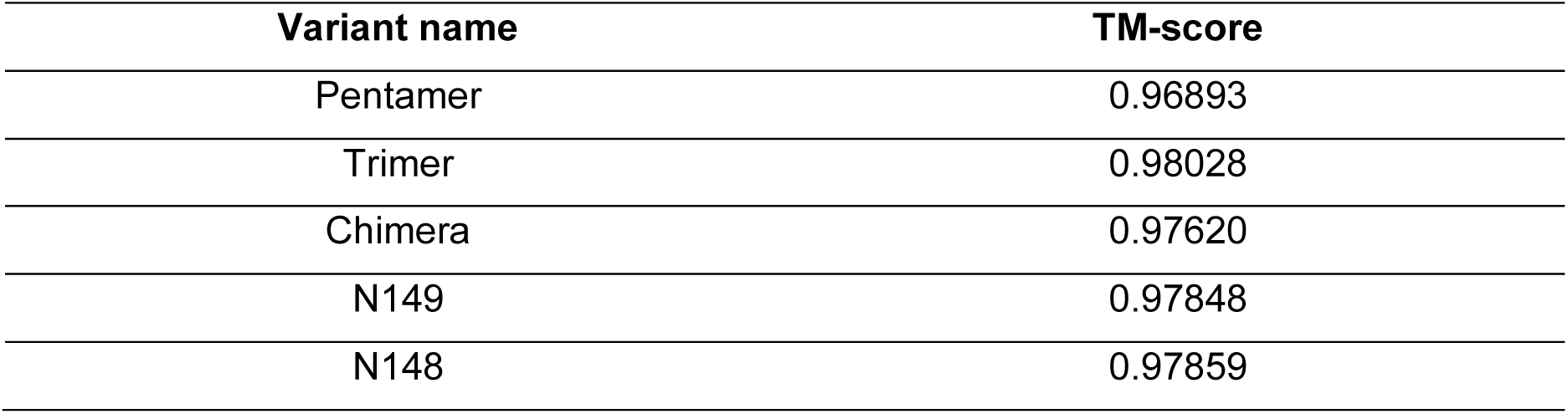
TM-score obtained from comparing AlphaFold predicted structures for redesigned variants to the wild-type AAV2 monomer structure.

| Variant name | TM-score |
| --- | --- |
| Pentamer | 0.96893 |
| Trimer | 0.98028 |
| Chimera | 0.97620 |
| N149 | 0.97848 |
| N148 | 0.97859 |

AAV2 monomer, with TM-scores above 0.9, where a value of 1 would indicate a perfect match between two structures.

Additionally, to visualise the structural overlap and differences between the AlphaFold predicted structures of the selected variants and the wild-type, the predicted variant monomer structures were superposed with the wild-type AAV2 monomer structure in Pymol (Figure 3). As expected, substantial overlap was observed across both loop and β-strand regions, indicating a high degree of structural alignment between the variants and the wild-type, and therefore a high confidence that the newly designed sequences would fold into the desired monomer structure.

**Figure 3.**
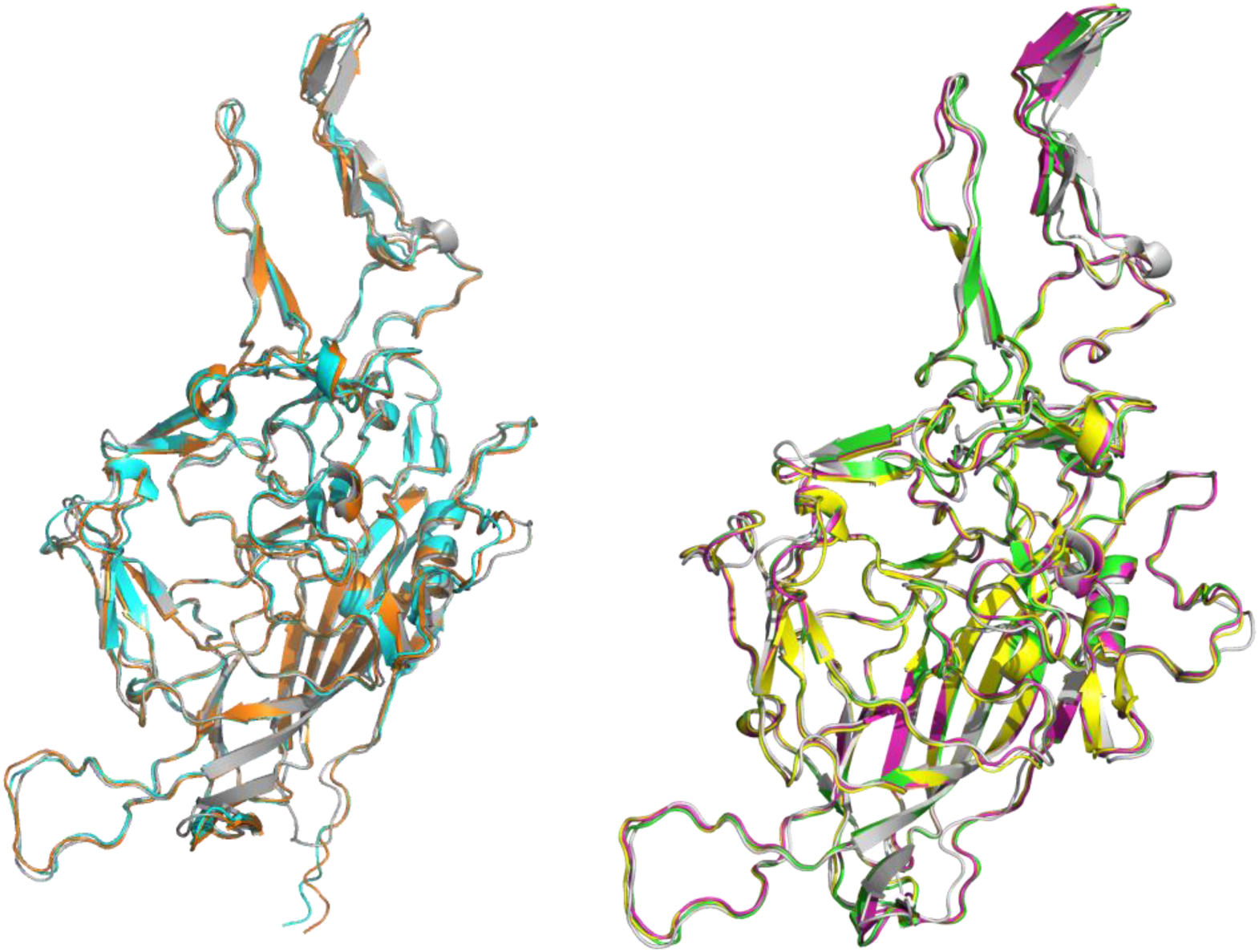
Predicted variant structures aligned with wild-type AAV2 monomer. A) Wild-type AAV2 monomer from PDB:1LP3 (grey) aligned against Pentamer consensus (cyan) and Trimer consensus (orange). B) Wild-type AAV2 monomer (grey) aligned against the Chimaera (yellow), N148 (magenta), and N149 (green). Structures of the variants were generated using AlphaFold 2 and aligned to WT in Pymol.

### ProteinMPNN and ancestral sequence reconstruction generated AAV2 variants could assemble into intact viral particle structures

The wild-type AAV2, two virtual ancestors, chimera, pentamer, and trimer redesigned AAVs, were each expressed in HEK293T cells and purified by ultracentrifugation to capture the combined layers containing both empty and full capsids. The capsids were then imaged by transmission electron microscopy (TEM) to determine whether intact capsids were formed. For all new designs, the TEM images showed pentagonal and hexagonal profiles of particles with diameters ranging from 17 to 25 nm (Figure 4), consistent with intact icosahedral wild-type AAV2 capsids.

**Figure 4.**
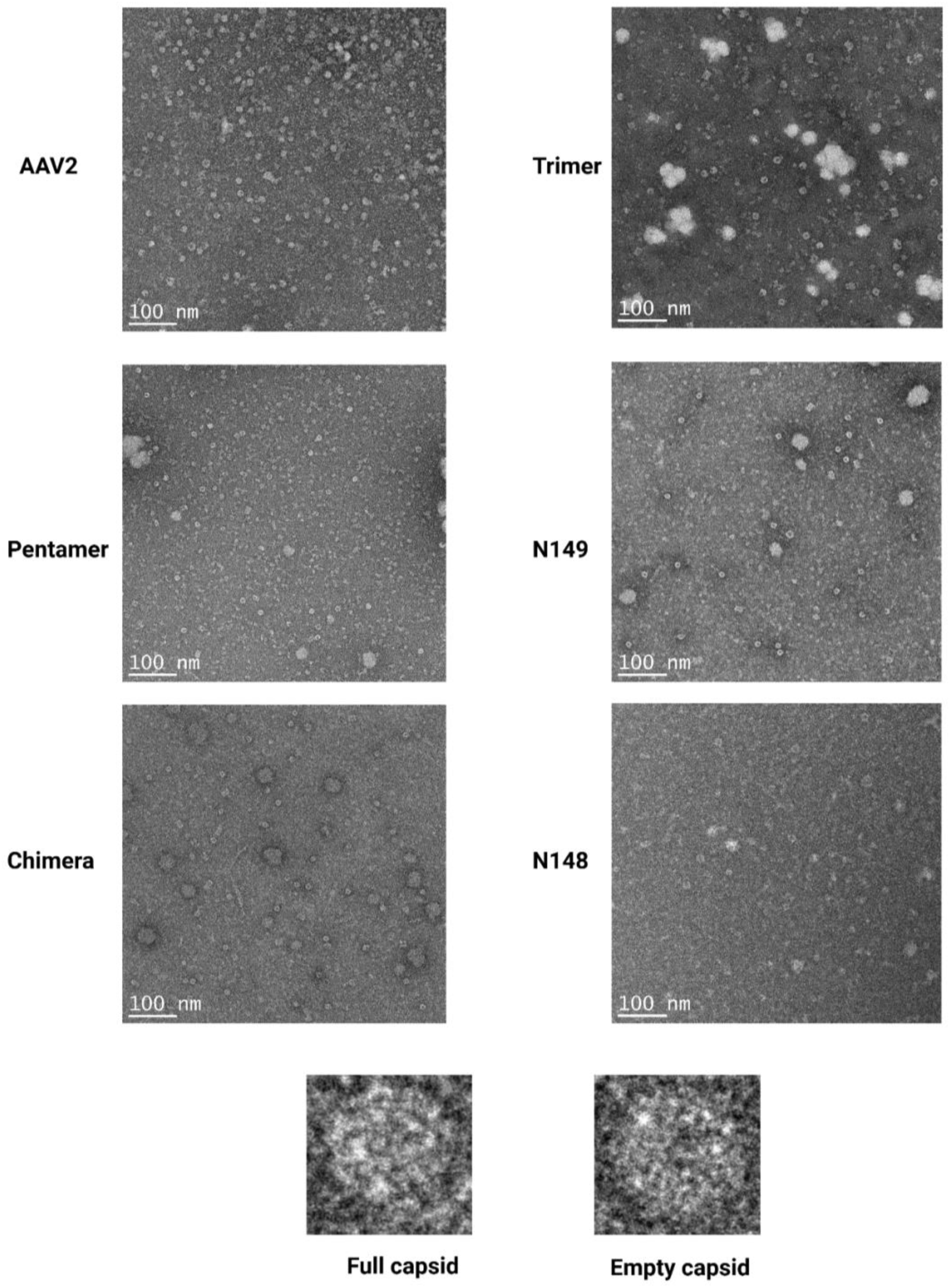
TEM images of selected re-designed AAV variants and wild-type AAV2, obtained under negative staining. In all images the expected “pentagonal” 2D representations of icosahedra are visible for both full and empty capsids of size range 17 to 25 nm. Many images also show evidence of larger non-uniform aggregate particles of 30-80 nm. Example zoomed-in images for individual full and empty capsids are shown at the bottom.

**Figure 5.**
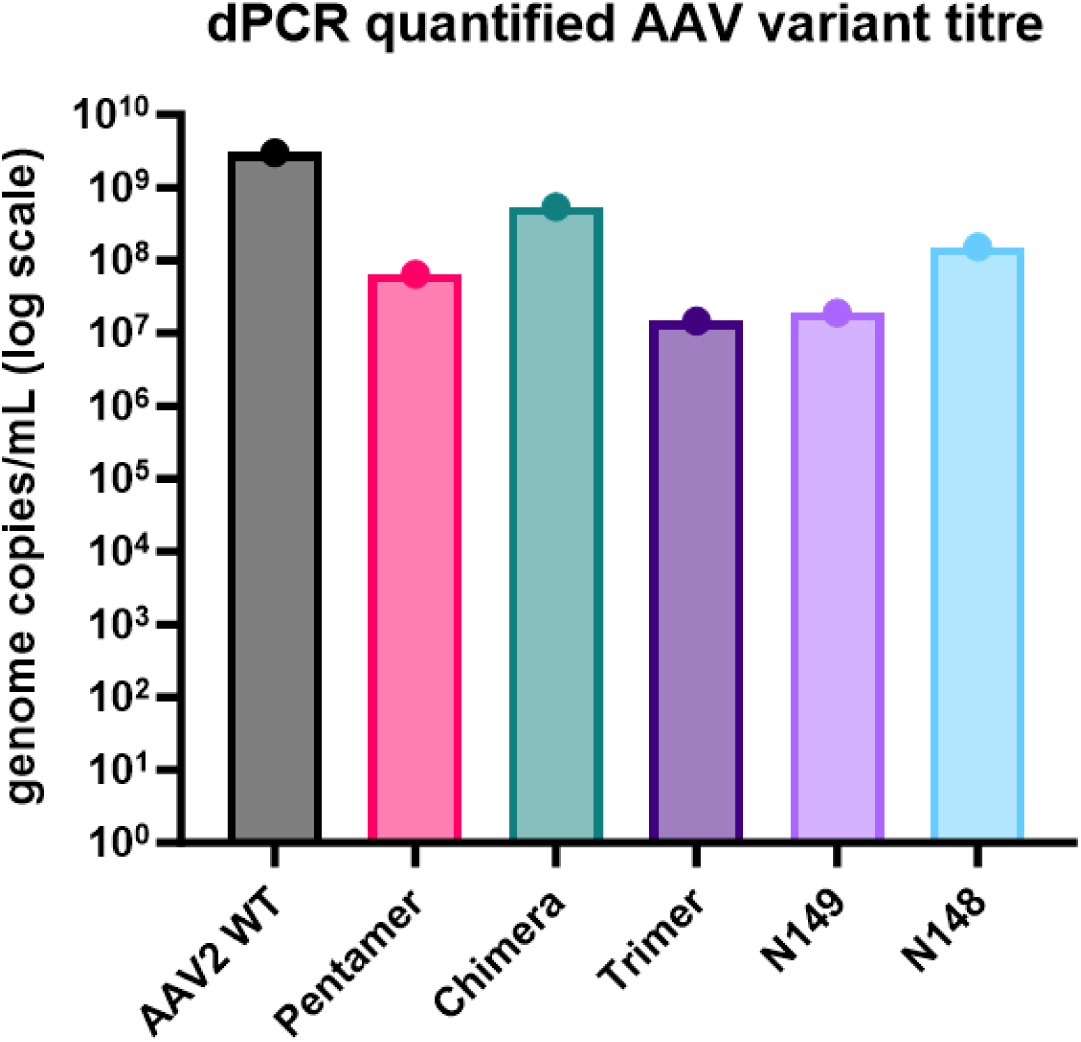
Genome titres determined by dPCR for AAV2 WT and redesigned variants. AAV samples were purified by ultracentrifugation from HEK293T cell cultures.

In all cases, both empty and full capsids were observed and were distinguishable due to the uranyl acetate staining used for TEM imaging (Figure 4). Capsids with dotted centres are considered empty or partially filled, given the staining of the inner capsid due to the absence of the transgene, whereas capsids that appear as homogenous spheres are considered to be filled capsids. Other than individual viral capsids, aggregates were also observed in some of the TEM images as clumps with irregular shapes. In the TEM images, varying amounts of aggregates are observed, suggesting different aggregation propensities of the variants.

The low concentration of virus in the samples made it difficult to quantify the relative amount of aggregates present in the samples, especially as aggregates can easily obscure AAV particles. However, based on the overall number of aggregates present across all negative-staining TEM images, Chimera, Pentamer and Trimer appear to have higher aggregation propensities.

The negative-staining TEM images clearly showed that the designed variants exhibited a similar morphology to wild-type AAV2. We deliberately did not separate empty from full capsids in the ultracentrifugation step so that we could assess the ratio of species formed in the cell culture step. Higher proportions of empty capsids were observed than filled capsids in all TEM images, indicating the low overall packaging efficiency of viral particles, even for the WT AAV2, using the HEK293T expression.

### ProteinMPNN and ancestral sequence reconstruction generated AAV2 variants were able to package DNA

To more accurately determine the packaging capacity of the novel variants and WT AAV2, the genome concentration was determined by droplet digital PCR (ddPCR). Figure 4 shows the titre in genome copies detected per mL of sample for all variants and wild type AAV2. The variants led to 5-200x lower titres than the wild type, ranging from 1.5×10^7^ to 5.4×10^8^ vg/ml (Table 2). The trimer and N149 variants, in particular, gave 200-fold lower titres than the wild-type AAV2. By contrast, the chimera variant stood out with a titre only 5x lower than for the wild-type AAV2.

**Table 2.** Summary of full capsid estimates from genome titres determined by dPCR and capsid titres determined by ELISA, for AAV2 WT and redesigned variants. AAV samples were purified by ultracentrifugation from HEK293T cell cultures without separating the empty from the full capsid layers.

| Variant name | Percentage of full capsids | Titre (vg/mL) | Total number of capsid (cp/ml) |
| --- | --- | --- | --- |
| AAV2 | 8.01% | $3.01 \times 10^9$ | $3.76 \times 10^{10}$ |
| Pentamer | 3.14% | $6.48 \times 10^7$ | $2.06 \times 10^9$ |
| Chimera | 20.47% | $5.44 \times 10^8$ | $2.66 \times 10^9$ |
| Trimer | 0.97% | $1.47 \times 10^7$ | $1.51 \times 10^9$ |
| N149 | 1.27% | $1.92 \times 10^7$ | $1.50 \times 10^9$ |
| N148 | 10.13% | $1.53 \times 10^8$ | $1.51 \times 10^9$ |

### The ProteinMPNN-guided designed variants show different levels of packaging efficiency

To determine the packaging efficiency of the designed variants, AAV capsid ELISA was performed to first quantify the total amount of AAV capsids present in each sample (Figure 6). Interestingly, the total capsid yields were broadly similar for the designed variants, and generally 10-20 times lower than for WT AAV2. The percentage of full capsids was calculated by dividing the viral genome/ml by the total capsid concentration. Out of all variants designed, the chimaera variant showed the highest percentage of the full capsid (20.5%), and this exceeded even that of wild-type AAV2 (8%), followed by variant N148 (5.1%). Whereas the percentages of full capsid for the trimer and N149 variants were the lowest, indicating poor packaging efficiency.

**Figure 6.**
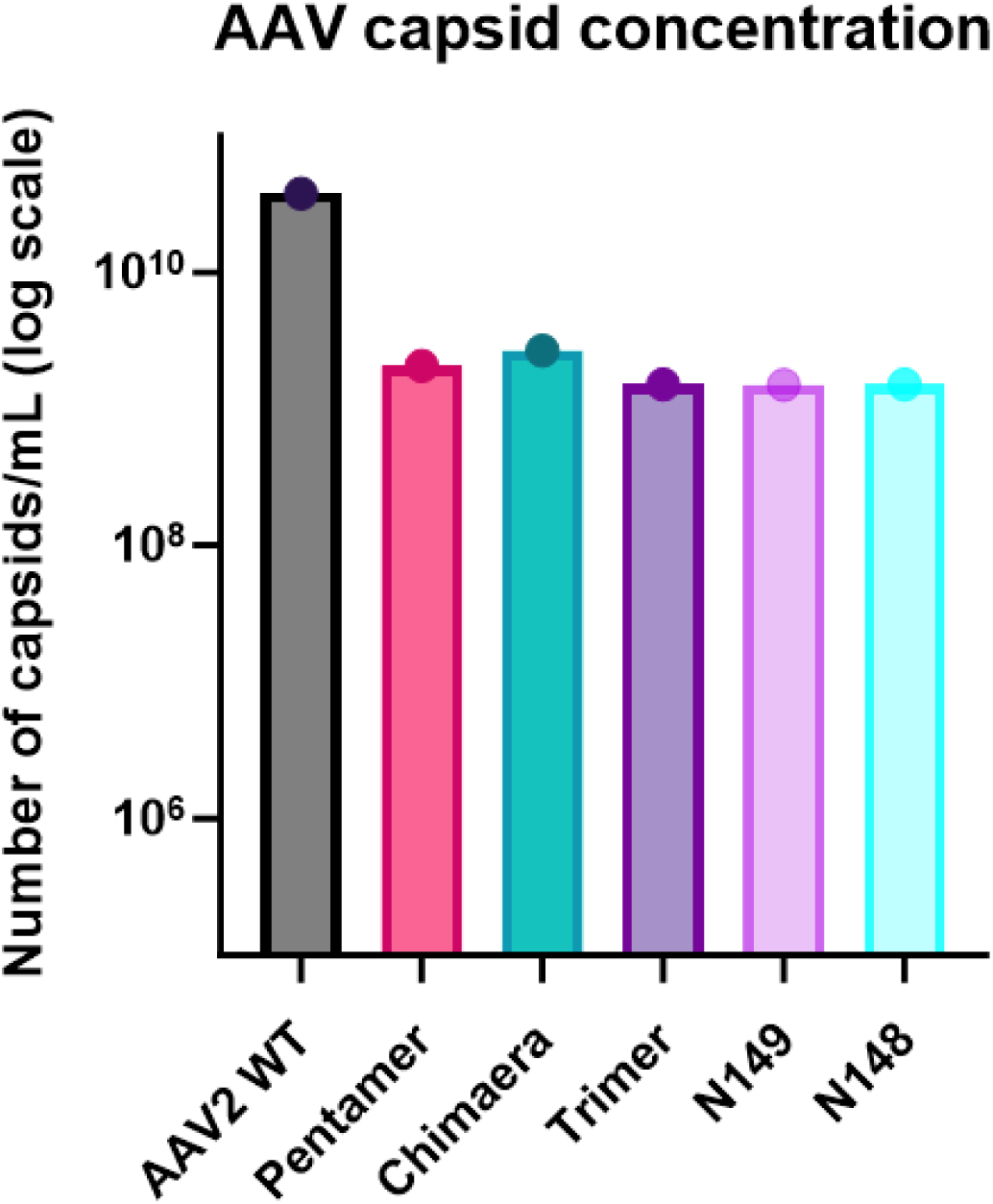
Capsid titres determined by ELISA for AAV2 WT and redesigned variants. AAV samples were purified by ultracentrifugation from HEK293T cell cultures.

These results indicate that all ProteinMPNN-based designs were all able to form intact capsids that gave yields of least 5% that of the WT AAV2. The key difference between capsid designs was in their packaging efficiency to yield full capsids.

### ProteinMPNN-guided designed variants retain functionality

The next question was whether the new capsid designs retained their ability to transduce HEK293 cells. Given that the designs retained the capsid surface sequence and redesigned only the core and interface structures of the capsid, the functionality of the vectors had the potential to be retained. Transduction efficiency was quantified using flow cytometry based on the GFP expression found in HEK293T cells and is shown in Figure 7. Indeed, it was found that all of the new capsid designs retained some level of transduction capability relative to WT AAV2, with the pentamer and trimer designs remarkably retaining similar transduction efficiencies to WT AAV2. The transduction efficiency of the chimera variant was 10x lower than for WT AAV2, having already accounted for the significantly higher packaging efficiency of this variant.

**Figure 7.**
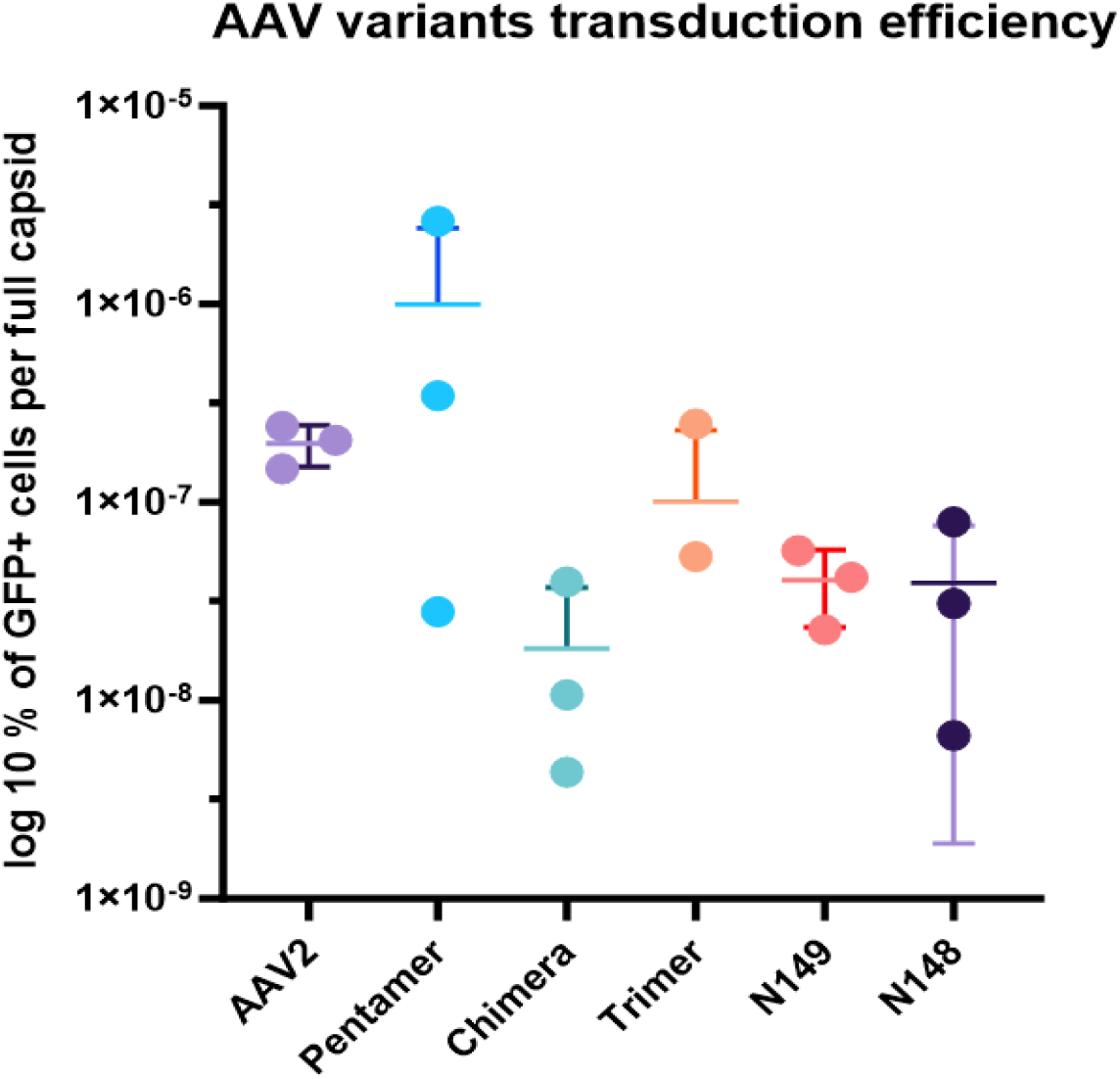
ProteinMPNN-guided AAV2 variant transduction efficiency normalised against the number of filled viral capsids added.

Overall, the similarity of transduction efficiency for all of the variants and wild type AAV2 indicated that this was governed primarily by the sequence and structural features of the solvent-exposed protein surface. It is well known that cell receptor binding is driven by surface feature recognition, such has heparin sulphate binding mediated by surface arginine/lysine residues mainly in the VIII loop at the trimer interface (Kern *et al*., 2003). However, it is still surprising that the transduction efficiencies of our variants were so similar given that the redesigns could easily have manifested differences through altered structural dynamics or changes in the stability and therefore disassembly of the capsid. By contrast, the retention of both the internally and externally solvent-exposed surface sequence and structure was not entirely sufficient to maintain the same level of DNA packaging efficiency. Indeed, redesign of the 5-fold axis channel through which DNA is thought to enter the capsids may be expected to alter capsid packaging efficiency. However, it is still perhaps surprising that the packaging efficiency only dropped by 16x in the worst variants (Trimer, N149) and by 5x in the pentamer. Clearly, the entry of DNA via the 5-fold axis channel can be controlled not only by the local mutations at the channel interfaces, but also by surrounding mutations that potentially modify the overall structure, dynamics or stability of the channel. This highlights the possibility of specifically engineering improved packaging efficiencies through mutations in and around the 5-fold axis.

Even more surprising was that the chimera design actually increased the packaging efficiency by approximately 2x despite retaining the pentamer interface mutations surrounding the 5-fold axis channel. A related surprising result from the design process was that the trimer and pentamer designs both retained the ability to form complete capsids despite allowing some unconstrained mutations at one of the subunit interfaces. Given that these two variants had low packaging efficiencies that were rescued (and improved over wild type) in the chimera, we can hypothesise that the lower performance of the trimer and pentamer designs was due to leakiness of the capsid to DNA via the non-optimal interface. In other words, the designs could still form intact capsids, but with one weakened interface. Therefore, the chimera design may have rescued the weakened interface to generate capsids that were better able to retain their DNA. An alternative mechanism through which our redesigned variants could have impacted DNA packaging is through mutations that also affect the interactions of VP3 surface residues with the Rep proteins responsible for genome packaging.

The attempts to re-optimise the core residues within the chimera design did not lead to any improvements and in fact, resulted in a loss of packaging efficiency again at least for one of the virtual ancestor variants selected. This optimisation in ProteinMPNN was carried out on the AlphaFold predicted monomer structure for the chimera. Therefore, any errors in the structure prediction would potentially compromise the ability to re-optimise within ProteinMPNN. By comparison, the trimer and pentamer redesigns used an x-ray crystal structure (PDB:1LP3) as the input, which is much less likely to contain structural errors. Alternatively, the re-optimisation could have led to increased capsid stability or altered dynamics that somehow hindered the packaging of DNA.

Another critical factor to consider is the potential influence of VP3 mutations upon interactions with the assembly-activating protein (AAP) during capsid assembly, and more critically, the impact of mutations within AAP itself. The gene for the 23 kDa (205 residue) AAP protein is nested into frame 2 of the *cap* gene, slightly preceding the start codon location of VP3 and overlapping with the first 177 residues of VP3. Therefore, 104 mutations in the VP3 region of the chimera may have impacted on AAP sequence and function.

AAP is involved in nucleolar localisation of VP from the cytoplasm/nucleus, as well as acting as a chaperone in capsid assembly. It is thought to interact with VP3 on the inner surface across the two-fold interface (Bennett, Mietzsch and Agbandje-Mckenna, 2017), while it is also known that the N-terminal region of AAP interacts with VP (Naumer *et al*., 2012).

As shown in Figures S4&5 (supplementary information), mutations were observed in all variants across the AAP regions that overlapped with the VP3 regions. All variants retained the first 42 residues identical to the wild-type AAP N-terminal hydrophobic region (HR) thought to interact with VP (Naumer *et al*., 2012), and also the first few residues of the conserved core. However, the remaining conserved core was heavily mutated in all variants, and all had a “TGA” stop codon after the conserved core but in the proline rich region (residue 64 or 66). It is therefore likely that the AAP in these variants had severely disrupted function. A previous study of AAP sequence deletions found that a complete truncation after the first 75 residues led to a knock-down of capsid titres to approximately 10% that of the wild-type (Tse *et al*., 2018). This is completely consistent with the 5% titres observed for our variants. All of the newly introduced stop codons were “TGA” codons, and so there is a possibility that some allowed a significant level of “readthrough” as is known for this stop codon in human cells (Trexler *et al*., 2023). However, the impact of the translated point mutations would still remain. In future we will consider addressing the re-encoding of VP3 for the new designs such that they incorporate fewer AAP mutations, or to co-express AAP from one of the production plasmids, and then re-evaluate their impact on vector production titres.

## Conclusion

This study demonstrates the powerful potential of generative AI tools, such as ProteinMPNN, in the rational design of AAV2 capsids. By allowing mutations in over half of the capsid residues—while preserving critical surface features—we successfully engineered variants with substantial sequence divergence (14–30%) that retained the ability to form stable, functional viral particles. Despite a general trend toward reduced titres in cell culture, most likely due to the simultaneous mutation of AAP, several designs exhibited enhanced functional properties. Notably, the “Pentamer” variant achieved superior transduction efficiency, while the “Chimera” variant demonstrated a 2.5-fold increase in packaging efficiency compared to wild-type AAV2. These findings demonstrate the feasibility of using AI-driven protein design to generate functional novel AAV capsids with significantly altered sequences to wild-type serotypes. Looking forward, this approach creates a platform from which we can begin to re-engineer these capsids even further to modify their tropism, to remove neutralising antibody epitopes, or to improve manufacturing titres. Additionally, the protein engineering approach directed to subunit interfaces and core residues could now be used to incorporate additional control elements or stability determinants at those interfaces, such as for pH-, ligand-or photo-selective disassembly.

## Acknowledgements

This research was conducted within and supported by the EPSRC Future Targeted Healthcare Manufacturing Hub at UCL (EP/P006485/1). Special thanks to Shu Chen at Birkbeck College London for support with Transmission Electron Microscopy (TEM) and Ben Jones at Queen Mary Blizard Institute, London for support with dPCR. The work was conceived and designed by ZJ and PAD. Materials were generated by ZJ and SL. Data was acquired by ZJ and SL. All analysis, interpretation and writing was carried out by ZJ, SL and PAD.

