## Supplementary information for "Fully functional AAV viral vectors with highly altered structural cores and subunit interfaces using ProteinMPNN"

**Figure S1. Sequences of AAV2 wild type and variants. Bold letters highlight the sites that were selected to mutate using ProteinMPNN (b-d), or the final mutations in new variants (e-i).**

a) Wildtype AAV2 (519 residues)

GADGVGNSSGNWHCDSTWMGDRVITTSTRTWALPTYNNHLYKQISSQSGASNDNHYFGYSTPWGYFDFNRFHCHFSPRDWQRL
INNNWGFRPKRLNFKLFNIQVKEVTQNDGTTTIANNLTSTVQVFTDSEYQLPYVLGSAHQGCLPPFPADVFMVPQYGYLTLNN
GSQAVGRSSFYCLEYFPSQMLRTGNNFTFSYTFEDVPFHSSYAHSQSLDRLMNPLIDQYLYYLSRTNTPSGTTTQSRLQFSQA
GASDIRDQSRNWLPGPCYRQQRVSKTSADNNNSEYSWTGATKYHLNGRDSLVNPGPAMASHKDDEEKFFPQSGVLIFGKQGSE
KTNVDIEKVMITDEEEIRTTNPVATEQYGSVSTNLQRGNRQAATADVNTQGVLPGMVWQDRDVYLQGPIWAKIPHTDGHFHPS
PLMGGFGLKHPPPQILIKNTPVPANPSTTFSAAKFASFITQYSTGQVSVEIEWELQKENSKRWNPEIQYTSNYNKSVNVDFTV
DTNGVYSEPRPIGTRYLTRNL

b) Pentamer interface (48 residues)

GADGVGNSSGNWHCDSTWMGDRVITTSTRTWALPTYNNHLYKQISSQSGASNDNHYFGYSTPWGYFDFNRFHCHFSPRDWQRL
INNNWGFRPKRLNFKLFNIQVKEVTQNDGTTTIANNLTSTVQVFTDSEYQLPYVLGSAHQGCLPPFPADVFMVPQYGYLTLNN
GSQAVGRSSFYCLEYFPSQMLRTGNNFTFSYTFEDVPFHSSYAHSQSLDRLMNPLIDQYLYYLSRTNTPSGTTTQSRLQFSQA
GASDIRDQSRNWLPGPCYRQQRVSKTSADNNNSEYSWTGATKYHLNGRDSLVNPGPAMASHKDDEEKFFPQSGVLIFGKQGSE
KTNVDIEKVMITDEEEIRTTNPVATEQYGSVSTNLQRGNRQAATADVNTQGVLPGMVWQDRDVYLQGPIWAKIPHTDGHFHPS
PLMGGFGLKHPPPQILIKNTPVPANPSTTFSAAKFASFITQYSTGQVSVEIEWELQKENSKRWNPEIQYTSNYNKSVNVDFTV
DTNGVYSEPRPIGTRYLTRNL

c) Trimer interface (124 residues)

GADGVGNSSGNWHCDSTWMGDRVITTSTRTWALPTYNNHLYKQISSQSGASNDNHYFGYSTPWGYFDFNRFHCHFSPRDWQRL
INNNWGFRPKRLNFKLFNIQVKEVTQNDGTTTIANNLTSTVQVFTDSEYQLPYVLGSAHQGCLPPFPADVFMVPQYGYLTLNN
GSQAVGRSSFYCLEYFPSQMLRTGNNFTFSYTFEDVPFHSSYAHSQSLDRLMNPLIDQYLYYLSRTNTPSGTTTQSRLQFSQA
GASDIRDQSRNWLPGPCYRQQRVSKTSADNNNSEYSWTGATKYHLNGRDSLVNPGPAMASHKDDEEKFFPQSGVLIFGKQGSE
KTNVDIEKVMITDEEEIRTTNPVATEQYGSVSTNLQRGNRQAATADVNTQGVLPGMVWQDRDVYLQGPIWAKIPHTDGHFHPS
PLMGGFGLKHPPPQILIKNTPVPANPSTTFSAAKFASFITQYSTGQVSVEIEWELQKENSKRWNPEIQYTSNYNKSVNVDFTV
DTNGVYSEPRPIGTRYLTRNL

d) Mutatable residues in pentamer and trimer designs (267 residues, 52%)

GADGVGNSSGNWHCDSTWMGDRVITTSTRTWALPTYNNHLYKQISSQSGASNDNHYFGYSTPWGYFDFNRFHCHFSPRDWQRL
INNNWGFRPKRLNFKLFNIQVKEVTQNDGTTTIANNLTSTVQVFTDSEYQLPYVLGSAHQGCLPPFPADVFMVPQYGYLTLNN
GSQAVGRSSFYCLEYFPSQMLRTGNNFTFSYTFEDVPFHSSYAHSQSLDRLMNPLIDQYLYYLSRTNTPSGTTTQSRLQFSQA
GASDIRDQSRNWLPGPCYRQQRVSKTSADNNNSEYSWTGATKYHLNGRDSLVNPGPAMASHKDDEEKFFPQSGVLIFGKQGSE
KTNVDIEKVMITDEEEIRTTNPVATEQYGSVSTNLQRGNRQAATADVNTQGVLPGMVWQDRDVYLQGPIWAKIPHTDGHFHPS
PLMGGFGLKHPPPQILIKNTPVPANPSTTFSAAKFASFITQYSTGQVSVEIEWELQKENSKRWNPEIQYTSNYNKSVNVDFTV
DTNGVYSEPRPIGTRYLTRNL

e) Pentamer subunit consensus (155 mutations, 30%)

GKDGIGNSSGNWHNDSTWNGDTVTLSRSWTFALPTYNNHKIKQVSSQSGVSNDNQYFGYTTPWGYLDWNRWSSFFSPRDWQRI
INNVLGFKPKRANFKIYNVQVFQVTQNDGTTTTTNLTTAEVAVFTDSEDQLPYVRGSLHQGCFPPYPGDVFLLPQYGYTTLWN
GSQALGRGSAYFPELLPSQVLRTGESYTFSYTFEDVPYIESYRYSQSIDRLLNPSVDQTTTVVSRTNAPSGTTTQSRTQYYQC
NASNWRDCSRNWLPGPIYRQQRLSKDSEDNNNSEYSWTGRATYHENGRDSPLNYGPAAATAKDDEEKYYPQNGVLVFGKQGSE
KTNADLEKLIIASYEELRPVNPYVYEQYGSVSTNVQRGNRQPGTRDVNTQGPLPGQIWADRPVYLDGPIWAKIPHTDGHSHPS
PRFGGFGLKHPFPQILAKITPVPFNPSTTYSPEKITSFITHYASGQVEVEVEVELQKENSKRWNPQRQLTSNYNKSNNLDFTY
DTNGNFSLPRMIGTRYVERNL

f) Trimer subunit consensus (71 mutations, 14%)

GSDGVGNSSGNWHCDSTWMGDTVITSRSRTWTLPTYNNHEYKQISSQSGASNDNQYYGYSTPWGYFDWNRWHCFFSPRDWQRL
INNVLGFRPKRLNFKIYNVQVKEVTQNDGTTTIANLTTATVQVFTDSEDQLPYVLGSLHQGCFPPFPADVFMLPQYGYLTLNN
GSQAIGRGSFYCLEYLPSQTLRTGDNYTFSYTFEDVPFHSSYVHSQSLDRSGNPLTDQYLYYLSRTNVPSGTTTQSRIQFSQP
DASDLRDQSRNWLPGPIYRQQRLSKDSSDNNNSEYSWTGRATYHLNGRDSALNPGPAMATAKDDEEKYFPQSGVLIFGKQGSE
KTNVDLEKVMITSEEELRPVNPVAYEQYGSVSTNLQRGNRQAGTEDVNTQGVLPGQVWQDRPVYLQGPIWAKIPHTDGHFHPS
PLMGGFGLKHPPPQILIKNTPVPANPSTTPSAAKFTSFITHYASGQVEVEVEWELQKENSKRWNPEIQLTSNYNKSVNEDFTR
DTNGVYSEPRPIGTRYLVRNL

g) Chimera (131 mutations, 25%) coloured by their location in VP3 (pentamer interface, trimer interface, core)

GKDGIGNSSGNWHNDSTWNGDTVTLSRSWTFALPTYNNHKIKQVSSQSGVSNDNQYFGYTTPWGYLDWNRWSSFFSPRDWQRI
INNVLGFKPKRANFKIYNVQVFQVTQNDGTTTTTNLTTAEVAVFTDSEDQLPYVRGSLHQGCFPPYPGDVFLLPQYGYTTLWN
GSQALGRGSAYFPELLPSQVLRTGESYTFSYTFEDVPFHSSYVHSQSLDRSGNPLTDQYLYYLSRTNVPSGTTTQSRIQFSQP
DASDLRDQSRNWLPGPIYRQQRLSKDSEDNNNSEYSWTGRATYHENGRDSPLNYGPAAATAKDDEEKYYPQNGVLVFGKQGSE
KTNADLEKLIIASYEELRPVNPVAYEQYGSVSTNLQRGNRQAGTEDVNTQGVLPGQVWQDRPVYLDGPIWAKIPHTDGHSHPS

P**RF**GGFGLKHP**F**PQIL**A**K**I**TPVP**F**NPSTT**Y**S**PE**K**IT**SFIT**H**Y**AS**GQV**E**VE**V**E**V**ELQKENSKRWNPEIQ**L**TSNYNKS**N**N**L**DFT**Y**
DTNG**NF**S**L**PR**M**IGTRYL**V**RNL

h) N149. (149 mutations, 28.8% mutated)

G**K**DG**I**GNSSGNWH**N**DSTW**N**GD**TIVLSR**T**W**T**FR**LPTYNNH**KI**KQ**V**SSQSG**V**SNDN**Q**YFGY**T**TPWGY**I**D**W**NR**WSAF**FSPRDWQR**I**
**V**NN**VV**G**YK**PKR**A**NFK**IYDV**QV**HQ**VTQNDGTTT**TT**N**LT**T**G**TV**E**VFTDSE**D**QLPY**IR**GS**L**HQG**V**LPP**NPN**D**LYTL**PQYGY**T**TL**W**N
GSQA**L**GR**G**S**A**Y**FP**E**LL**PSQ**V**LRTG**ESY**T**Y**SYTFEDVPFHSSY**V**HSQSLDR**SG**NPL**T**DQYLYYLSRTN**V**PSGTTTQSR**I**QFSQ**P**
**D**ASDLRDQSRNWLPGP**K**YR**D**QR**L**SK**D**S**S**DNNNSEYSWTGA**NT**YH**E**NGRDS**PM**N**L**GPA**T**ASHKDDEEK**YYG**Q**QSQ**L**VY**GKQGSE
KTN**A**D**L**EK**LL**I**E**D**T**EE**L**R**PV**NPVA**Y**EQYGSVSTNLQRGNRQA**G**T**E**DVNTQGVLPG**Q**VWQDR**PDT**L**D**GPIW**T**KIPHTDGH**S**HPS
P**RF**GGFGLKHP**F**P**L**IL**ARI**TPVP**F**NPSTT**Y**S**PE**K**IT**SFITQY**A**TG**NIEI**E**L**E**V**ELQKENSKRWNPEIQ**L**TSNYNKS**N**N**L**DFT**Y**
DTNG**NF**S**I**PR**LMSD**RYL**V**RNL

i) N148 (150 mutations, 29% mutated)

G**K**DG**I**GNSSGNWH**N**DSTW**N**GD**TIVLKR**T**W**T**FR**LPTYNNH**KI**KQ**V**SSQSG**V**SNDN**Q**YFGY**T**TPW**A**Y**I**D**W**NR**WSAF**FSPRDWQR**I**
**V**NN**VV**G**YK**PKR**A**NFK**IYDV**QV**YQ**VTQNDGTTT**TT**N**LT**T**G**TV**E**VFTDSE**D**QLPY**IR**GS**L**HQG**V**LPP**N**P**N**D**LYTL**PQYGY**T**TL**W**N
GSQA**L**GR**G**S**A**Y**FP**E**LL**PSQ**V**LRTG**ESY**T**Y**SYTFEDVPFHSSY**V**HSQSLDR**SG**NPL**T**DQYLYYLSRTN**V**PSGTTTQSR**I**QFSQ**P**
**D**ASD**L**RDQSRNWLPGP**K**YR**D**QR**L**SK**D**S**S**DNNNSEYSWTGA**NT**YH**E**NGRDS**PM**N**L**GPA**T**ASHKDDEEK**YYG**Q**QSQRV**FGKQGSE
KTN**A**D**L**EK**LL**I**E**D**T**EE**L**R**PV**NPVA**Y**EQYGSVSTNLQRGNRQA**G**T**E**DVNTQGVLPG**Q**VWQDR**PDT**L**D**GPIW**S**KIPHTDGH**S**HPS
P**RF**GGFGLKHP**F**P**L**IL**ARI**TPVP**F**NPSTT**Y**S**PE**K**IT**SFITQY**A**TG**NIEI**E**L**E**VA**LQKENSKRWNPEIQ**L**TSNYNKS**N**N**L**DFT**Y**
DTNG**NF**S**I**PR**IMSD**RYL**V**RNL

**Figure S2 Phylogenetic tree showing the position of N149 and N148 visualised using iTOL.**

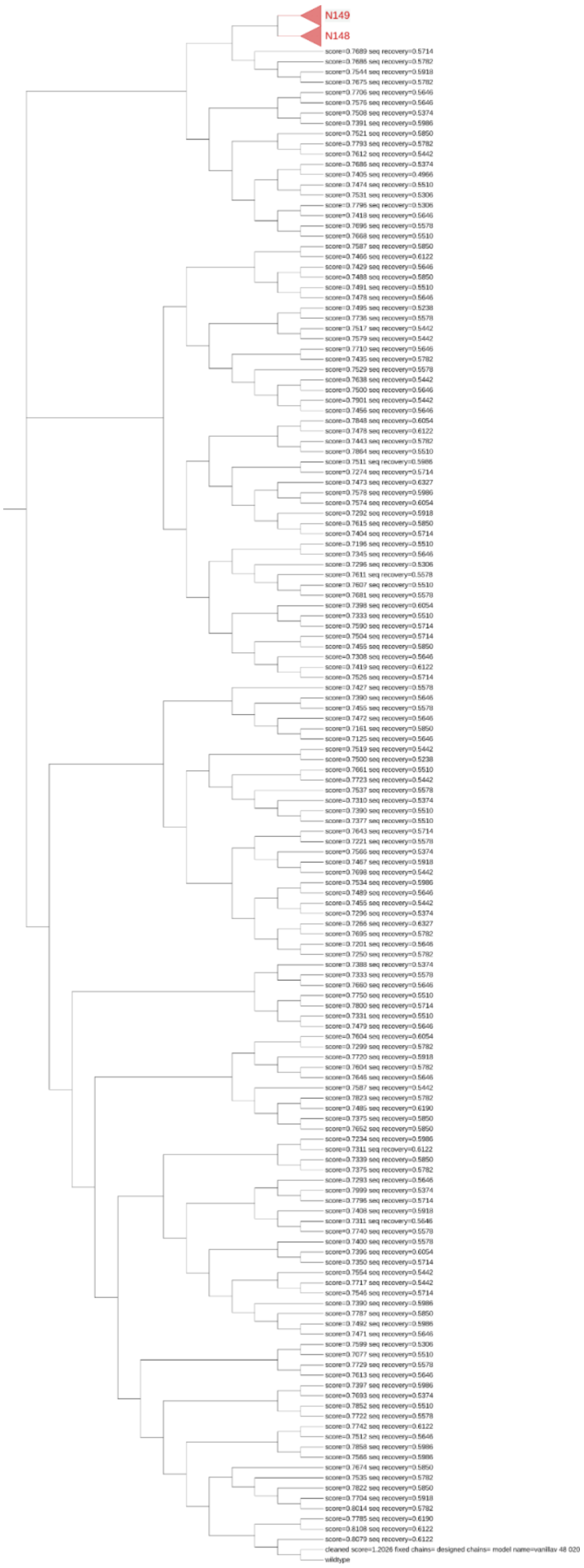

**Figure S3. Solubility scores predicted by Camsol (Sormanni, Aprile and Vendruscolo, 2015) for** A) 150 "virtual" ancestor chimera sequences, and B) 150 original ProteinMPNN redesigned chimera sequences. The new AI-generated sequences are shown in blue, while the original parent wild-type AAV2 sequence is shown as a red dot and the chimera consensus sequence is shown as a magenta dot.

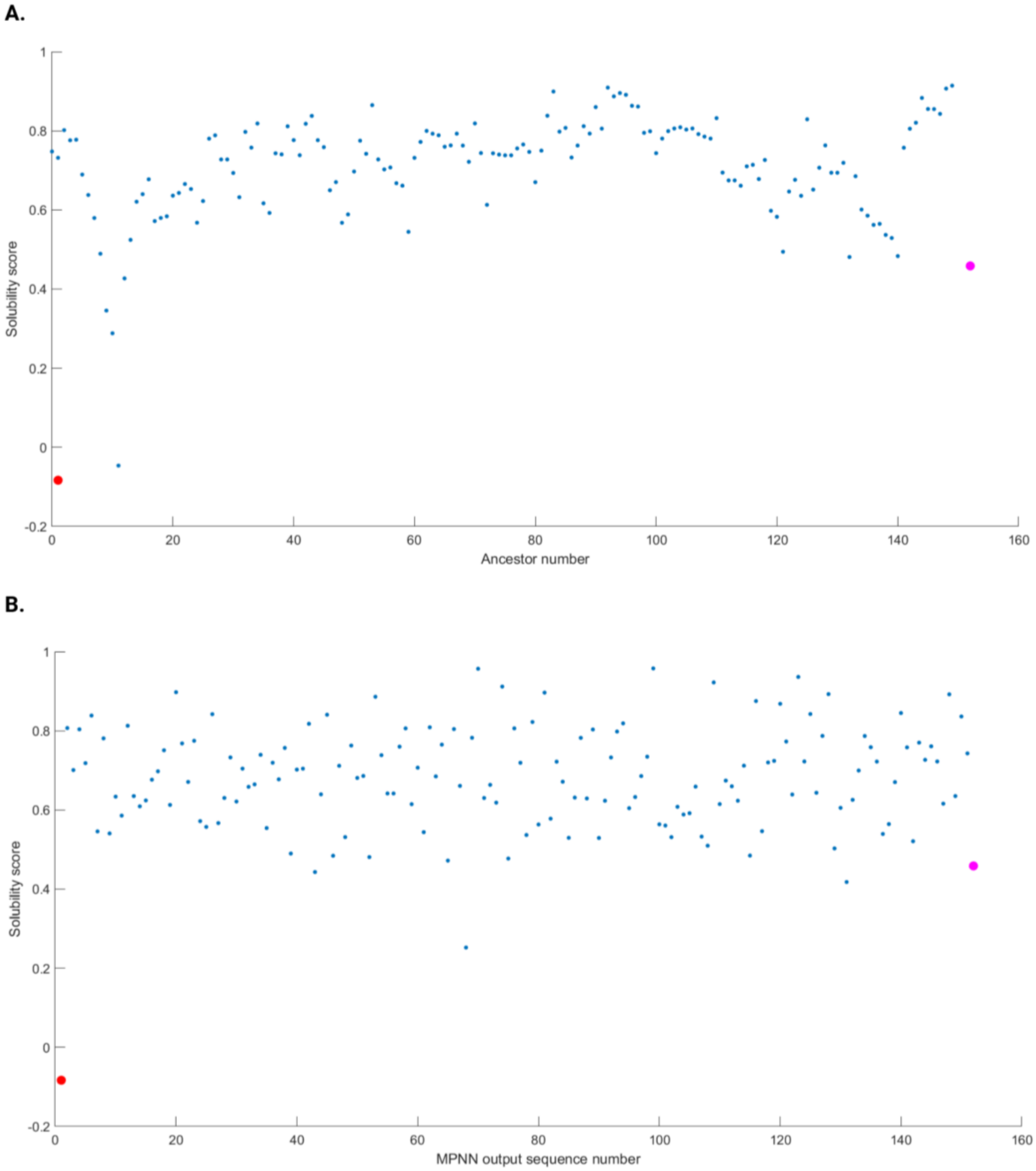

**Figure S4 AAP region DNA sequence alignments for the variants and wild-type. Alignments were generated using the SnapGene MUSCLE alignment algorithm.**

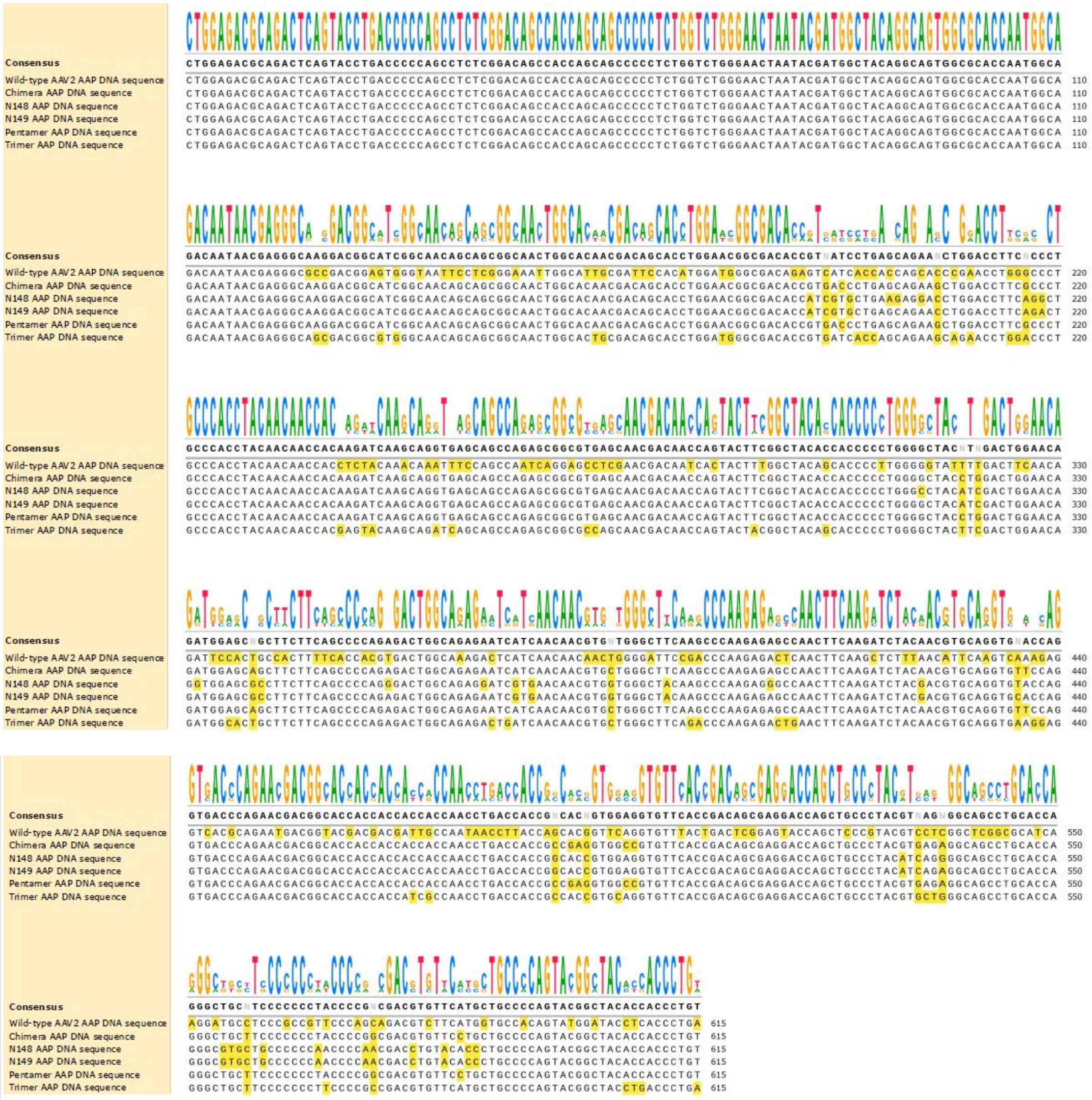

**Figure S5 AAP region protein sequence alignments for the variants and wild-type.** Alignments were generated in SnapGene using the CLUSTAL OMEGA algorithm. Stop codons are represented by * in the protein sequences.

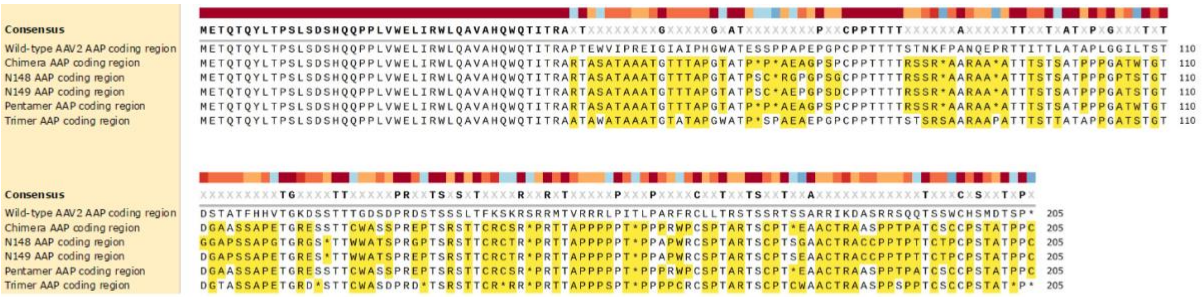
